# The Mechanism of Vesicle Solubilization by the Detergent Sodium Dodecyl Sulfate

**DOI:** 10.1101/2020.06.05.133868

**Authors:** José Juan-Colás, Lara Dresser, Katie Morris, Hugo Lagadou, Rebecca H. Ward, Amy Burns, Steve Tear, Steven Johnson, Mark C. Leake, Steven D. Quinn

## Abstract

Membrane solubilization by sodium dodecyl sulfate (SDS) is indispensable for many established biotechnological applications, including viral inactivation and protein extraction. Although the ensemble thermodynamics have been thoroughly explored, the underlying molecular dynamics have remained inaccessible, owing to major limitations of traditional measurement tools. Here, we integrate multiple advanced biophysical approaches to gain multi-angle insight into the time-dependence and fundamental kinetic steps associated with the solubilization of single sub-micron sized vesicles in response to SDS. We find that the accumulation of SDS molecules on in-tact vesicles triggers biphasic solubilization kinetics comprising an initial vesicle expansion event followed by rapid lipid loss and micellization. Our findings support a general mechanism of detergent-induced membrane solubilization and we expect the framework of correlative biophysical technologies presented here will form a general platform for elucidating the complex kinetics of membrane perturbation induced by a wide variety of surfactants and disrupting agents.

## Introduction

The solubilization of lipid membranes by the anionic detergent sodium dodecyl sulfate (SDS) has found vast utility across the pharmaceutical and biological sectors with far-reaching applications including rapid cell lysis, protein extraction and viral inactivation^[1-3]^. However, despite decades of empirical use, accessing the molecular mechanisms through which SDS leads to membrane solubilization has remained a major experimental challenge, owing to the shortcomings of technologies which rely on averaging over the entire process^[4, 5]^.

Initial steady-state thermodynamic experiments reported the partitioning of SDS into lipid membranes, illuminating several key factors, including the detergent: lipid molar ratio, surface pressure, electrostatic forces and temperature, that modulate the initial interaction^[6, 7]^. These experiments also indicated that the SDS-membrane partition coefficient is similar to those obtained for non-ionic detergents of identical chain lengths^[7]^. The kinetics of the following disruption process have also been investigated at the ensemble level using fluorescence strategies, whereby the emission intensity of small fluorescent molecules encapsulated into vesicles increased due to dye diffusion across the perturbed bilayer^[8]^. Here, vesicle leakage was characterized by a single exponential rate of release with the rate constant increasing with SDS concentration according to a power-law dependence. Differential scanning calorimetry has also been employed to examine the temperature-dependence and formation of mixed detergent-lipid micelles at the end-point of the solubilization process, revealing that membrane degradation is related to hydrophobic interactions between the SDS molecule and lipids^[9]^.

While it is abundantly clear from such studies that SDS influences the membrane structure, analyses have been constrained to the identification of structural changes in the ensemble ^[6, 10, 11]^, and while such data reveals valuable information about the mean average state of the interaction, dynamic heterogeneity within the sample, in addition to the presence of rapid transient intermediate molecular-level events remains obscure. Nevertheless, the combination of ensemble-based approaches have broadly alluded to a three-step model of solubilization^[12]^. In step one, detergent monomers saturate the membrane, leading to the co-existence of mixed detergent-lipid micelles within the intact membrane in step two. In step three, solubilization is achieved via the fragmentation and release of mixed detergent-lipid micelles into the solution. However, whether these steps take place sequentially or if they are interconnected remains an open question, and uncertainties left on the molecular mechanisms are due to a lack of techniques that can capture the time-dependent dynamics of the interaction across the entire solubilization window.

While protocols from the Cryo-TEM and NMR communities have provided important structural insights into membrane conformations in response to SDS^[13, 14]^, neither report on the underlying time-dependent dynamics. Conversely, isothermal calorimetry and turbidity measurements have revealed the relative detergent: lipid stoichiometry ratios required to achieve complete solubilization, but neither quantify changes within the membrane architecture^[15-17]^.

Molecular dynamics simulations have gone some way to bridging this gap by enabling discrete steps of the initial interaction, including structural reorganizations in the membrane^[18, 19]^ and micellization^[20]^, to be followed over several tens of nanoseconds. More generally, these studies also support an interaction between the surfactant and lipid head groups which leads to the penetration of SDS into the membrane, in turn increasing the bilayer thickness and lipid tail order. Hydrophilic interactions between the sulfate and phosphocholine groups, and hydrophobic interactions between hydrocarbon surfactant and lipid chains also gives rise to a tight packing density and ordered chain alignment. Additionally, optical microscopy based on the phase contrast and fluorescence imaging of giant unilamellar vesicles also indicate changes in membrane curvature after SDS incorporation, closely followed by the stress-induced formation of macropores and fragmentation^[21]^. At SDS concentrations lower than the critical micelle concentration (CMC), transmission microscopy experiments also indicate the homogeneous distribution of SDS monomers into the membrane, followed by the gradual formation of mixed micelles and at concentrations around the CMC, local instabilities in the vesicle architecture are introduced across the structure^[22]^. Unfortunately, these microscopy tools are limited to a spatial resolution of ∼250 nm and cannot access the solubilization dynamics of vesicles which are smaller than the optical diffraction limit and which have high radii of curvature. Moreover, conventional microscopy tools can only measure macroscopic changes in membrane structure and packing density, and provide little structural and mass information on the nanoscale, motivating the need for the multi-disciplinary approach outlined here to gain molecular-level insight into this important biological problem.

To overcome the hurdle posed by the optical diffraction limit, we recently reported a structural imaging method based on single-vesicle Förster resonance energy transfer (svFRET) to study rapid organizational changes in sub-micron sized vesicles in response to the non-ionic detergent Triton-X 100^[23]^. A key benefit of the svFRET approach is the ability to access time-dependent solubilization kinetics from single immobilized vesicles with millisecond time resolution, bypassing the ensemble average.

Here, we combine the svFRET technology with steady-state fluorescence spectroscopy, dynamic light scattering (DLS), liquid-based atomic force microscopy (AFM) and quartz-crystal microbalance with dissipation monitoring (QCM-D), to reveal the fundamental solubilization steps and kinetics associated with sub-micron sized and highly curved vesicles in response to SDS, without interference from vesicle fusion. We unveil the solubilization mechanism as a three step process in which detergent accumulation on the LUV surface precedes biphasic solubilization kinetics consisting of an initial expansion of the vesicle, followed by rapid lipid loss. The capability of discriminating between solubilization steps and kinetic parameters is a novel finding only afforded by the implementation of these correlated approaches, and we expect the presented strategy should assist in the effective and efficient fine-tuning of detergent-induced solubilization protocols, and for exposing the multifaceted dynamics of membrane perturbation caused by a broad range of molecular disruptors.

## Results and Discussion

LUVs composed of 98.8% 1,2-dimyristoyl-sn-glycero-3-phosphocholine (DMPC), 1% 1,2-dioleoyl-sn-glycero-3-phosphoethanolamine-N-(biotinyl) (Biotin-PE), 0.1 % Dil (svFRET donor) and 0.1 % DiD (svFRET acceptor) were prepared as detailed in the **Methods** section and are schematically illustrated in **Figure S1**. DMPC is a synthetic phospholipid used widely for the preparation of vesicles and supported lipid bilayers^[24]^, and is used here to provide a structural framework. Dil and DiD are lipophilic cyanine derivatives and when a molar ratio per vesicle of 1:1 was used, the average spatial separation between them was close to their Förster radius of 53 Å, corresponding to an average apparent FRET efficiency (E_FRET_) of ∼0.5. As we previously reported^[23]^, this enables nanoscopic changes in the average inter-dye distance to be measured by observable changes in E_FRET_ in either direction.

The formation of ∼200 nm-sized LUVs was confirmed by DLS (**Figure S2**), and their steady-state fluorescence emission was recorded as a first step to characterize their interaction with SDS. We note that the hydrodynamic radii and ensemble E_FRET_ values obtained from the LUVs remained largely invariant over the course of several weeks, pointing towards their long-term stability under the buffer conditions used (**Figure S3**). As the concentration of SDS was increased towards the CMC, a progressive increase in Dil emission, concurrent with a progressive decrease in DiD emission was observed, giving rise to an overall decay in E_FRET_ (**Figure 1a**), from an initial value of 0.50 ± 0.02 (± SEM) in the absence of SDS to 0.09 ± 0.02 in the presence of 4 mM. A Hill model applied to the E_FRET_ data indicates a half-maximal concentration constant of 0.92 ± 0.16 mM with a Hill coefficient of 1.25 ± 0.20. The time-dependent increase in Dil emission, immediately after injection of SDS followed bi-exponential kinetics, where the amplitude weighted average time-constant progressively decreased from 28.2 ± 7.6 s in the presence of 1 mM SDS to 4.51 ± 0.48 s at 4 mM (**Figure S4** and **Table S1**). Taken together, these data suggest that the addition of SDS at concentrations approaching the CMC introduces structural changes in the vesicle architecture, composition, or both, that results in a ∼47 % increase in the average spatial separation distance between the fluorophores.

**Figure 1.**
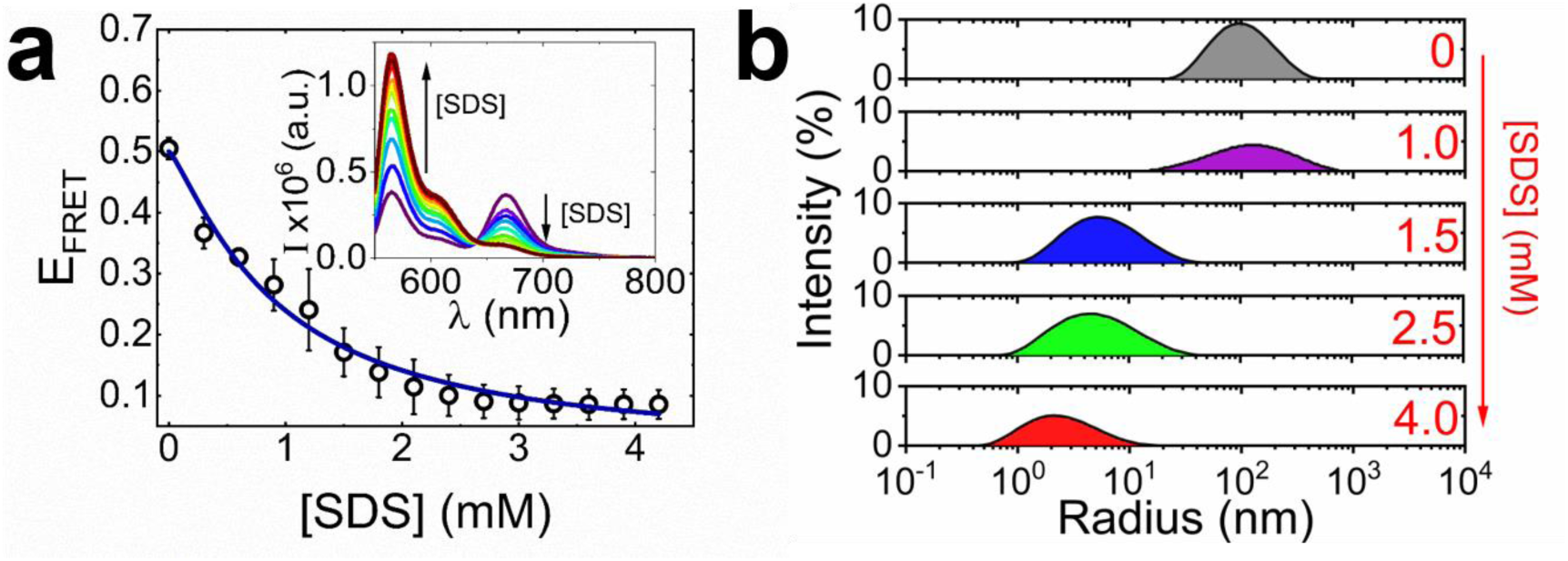
SDS vesicle interactions reported using ensemble FRET spectroscopy and dynamic light scattering. (a) FRET efficiency of Dil/DiD labeled vesicles versus SDS. The solid line represents a Hill model fit. Inset: the corresponding variation in fluorescence spectra. (b) DLS size distributions observed from the free diffusion of vesicles in solution in the absence and presence of 1 mM, 1.5 mM, 2.5 mM and 4 mM SDS.

Having established ensemble FRET as an optical sensor of changes in dye separation distance as a function of SDS, DLS was next used to probe the hydrodynamic radii of LUVs in response to SDS in solution. The high sensitivity of DLS to the diffusion of nanomolar concentrations of vesicles makes it an attractive technique for accessing structural characteristics along the solubilization pathway^[25, 26]^. Here, the autocorrelation function obtained from the intensity of light scattered by the LUVs yields their translational diffusion coefficient which can be used to extrapolate their hydrodynamic radius (r_H_) via the Stokes-Einstein equation. In the absence of SDS, the LUVs were distributed around a peak r_H_ of 97 ± 6 nm (**Figure 1b**). However, the peak r_H_ associated with the LUVs in the presence of 1 mM SDS was found to increase by 25 %, and the full-width at half-maximum (FWHM) of the distribution broadened two-fold. At higher SDS concentrations, a progressive decrease in peak r_H_ towards several nanometers was observed, which we attributed to the formation of micelles in solution. The increase in peak r_H_ observed at relatively low SDS concentrations, points towards an increase in mean LUV surface area prior to micellization and was assigned to LUV swelling, fusion, or the combination of both in solution. As recently demonstrated by molecular dynamics simulations^[27]^, the high membrane curvature associated with LUVs regulates recruitment and may in this case facilitate the incorporation of SDS monomers into the bilayer, even at concentrations lower than the reported CMC.

To rule out the possibility of fusion and investigate each step of the solubilization process in more detail, LUVs containing a low percentage of Biotin-PE were immobilized onto an Avidin-coated surface and imaged by liquid-based atomic force microscopy in the absence and presence of SDS. As shown in **Figure S5**, biotinylated vesicles were tethered to Avidin which was in turn coupled to biotinylated bovine serum albumin adsorbed onto a glass substrate. In the absence of SDS, the LUVs appeared spherical in nature (**Figure 2a**) with a mean caliper diameter, defined as the average LUV width at half maximum of approximately 44 ± 1 nm (±SEM, N=50) (**Figure 2b**). In the presence of low concentrations of SDS, namely 0.1 mM and 0.5 mM SDS, the mean caliper diameters were found to be approximately 51 ± 2 nm (N=50) and 57 ± 2 nm (N=50), corresponding to a size increase of 16.1 ± 0.5 % and 28.3 ± 0.9 %, respectively (**Figure 2b**). In turn, these data suggest an interaction between SDS and the vesicles that leads to LUV expansion. We note that the measured LUV heights ranged from approximately 18 – 30 nm, similar to those previously reported for extracellular vesicles of comparable size. The observed reduction in LUV height relative to the diameter measured by DLS was assigned to elastic deformation of the structures induced by mechanical indentation exerted by the AFM tip as previously reported^[28-30]^. Here, elastic deformation occurs at the onset of interaction and a tip indentation of ∼10 nm is typical with applied forces of ∼2 nN, comparable to those used in this work^[31]^. Nevertheless, the measurable expansion of mean caliper diameters in the presence of SDS points towards a mechanism of interaction involving a substantial LUV structural change that precedes complete solubilization.

**Figure 2.**
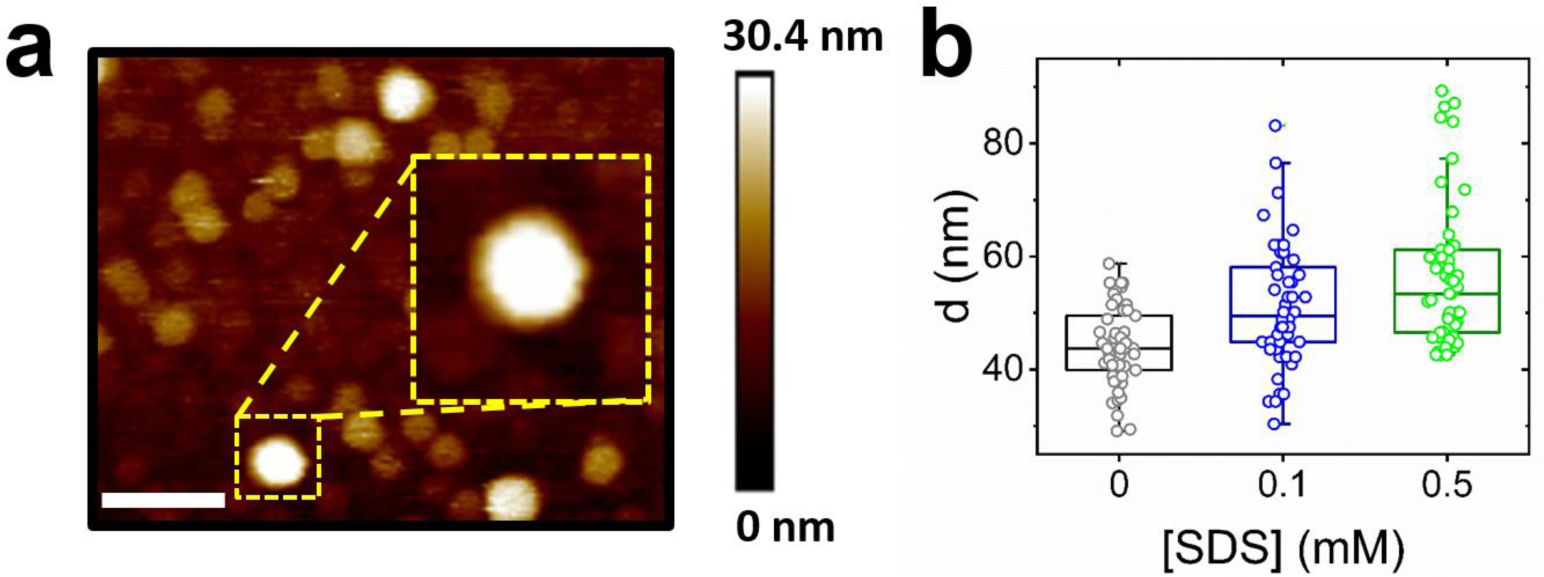
Visualization of SDS-induced vesicle expansion by liquid AFM. (a) Representative image of LUVs pre-immobilized onto a glass substrate using a BSA-biotin-Avidin immobilization scheme. The scale bar indicates 150 nm. (b) Comparative box plots, first quartile, median, third quartile and standard deviation (error bars) are shown summarizing the relative variation in mean caliper diameter, d, in the absence (grey, N = 50) and presence of 0.1 mM (blue, N = 50) and 0.5 mM (green, N = 50) SDS.

To investigate this observation further, and to extract the kinetic rates of the swelling event, single immobilized vesicles were imaged via objective-based total internal reflection fluorescence (TIRF) microscopy and the E_FRET_ response recorded with a time-resolution of 50 ms as SDS was flushed across the sample. To ensure fluorescence signals originated from the Dil and DiD labelled LUVs, varying concentrations of vesicles were added to the substrate, resulting in a corresponding change in the number of fluorescent foci per field-of-view (**Figure S6**). To optimize the experimental conditions necessary for minimizing photobleaching, fluorescence emission from single LUVs in the absence of SDS were first recorded as a function of excitation intensity. Here, Dil and DiD labelled LUVs were imaged using 532 nm excitation powers of 5 mW, 1 mW and 16 μW, as measured at the back aperture of the TIRF objective lens. When the excitation power was 5 mW, the fluorescence trajectories displayed bi-exponential behavior (R^2^ = 0.97) with decay time constants of 1.6 ± 0.1 s and 10.8 ± 0.6 s, respectively (**Figure S7**). At 1 mW, the fluorescence decays were dominated by longer-lived components and an average decay time of 23.7 ± 0.3 s was observed. At 16 μW, however, the trajectories were photostable across the duration of the measurement time window (**Figure S7**). Having established that an excitation intensity of 16 μW was necessary for long-term photostability, LUV perturbation induced by SDS was then detected through changes in the apparent FRET efficiency. Conformational changes were first determined via histograms of the svFRET efficiency as a function of SDS concentration (**Figure 3a**). The E_FRET_ histogram for LUVs in the absence of SDS showed a single Gaussian peak at 0.39 ± 0.1 (FWHM = 0.10), characteristic of an intact LUV in which the average fluorophore separation is 5.7 ± 1.5 nm. After incubation of the immobilized LUVs with intermediate SDS concentrations, a progressive decrease in the peak position towards 0.19 ± 0.06 (FWHM = 0.13) at 0.20 mM SDS was observed, indicating an 18 % increase in the mean inter-dye distance. In the presence of 1.0 mM SDS, a doublet distribution, reflecting heterogeneity in the immobilized LUVs, and assigned to partial lysis was observed, with the main peak position shifting further to 0.14 ± 0.02 and corresponding to an overall 26 % increase in dye separation. Taken together, the overall reduction in the svFRET signal, concurrent an increase in the DiL-DiD separation distance, agrees well the ensemble FRET measurements and AFM data. The fraction of intact LUVs estimated from the relative area under the histogram in the absence of SDS also progressively decreased, with negligible levels of intact and immobilized LUVs present when 10 mM SDS was applied. Next, to extract the kinetic details of the interaction, fluorescence emission trajectories obtained from single LUVs were recorded over time as SDS was injected into a flow-cell containing pre-immobilized LUVs. In the absence of SDS, emission trajectories obtained from Dil and DiD were invariant with no variation in the apparent FRET efficiency observed. However, as SDS was injected into the flowcell, deformation of single LUVs was then measured via anti-correlations in the Dil and DiD signals and thus quantifiable changes in the apparent FRET efficiency (**Figure 3b and Figure S8**). In the absence of SDS, E_FRET_ from single vesicles was found to be invariant with a value of 0.32 ± 0.01. Injection of 10 mM SDS then induced variations in E_FRET_ and the total intensity over very different timescales. Immediately after injection, the FRET efficiency decreased to 0.06 ± 0.01 over 8.5 seconds **(Figures 3b and 3c**) with most of this change occurring within the first 5.5 seconds, pointing towards an initial 18 % increase in the average separation between the dyes. Assuming spherical vesicles, this increase scales directly with the vesicle radius and thus agrees well with the swelling observed by AFM. Within this time window, the total intensity associated with each vesicle remained invariant, however, at longer timescales (> 23 s) E_FRET_ remained constant at a lower value of 0.06 ± 0.01, whereas the total intensity decreased rapidly to only a few percent of its initial value.

**Figure 3.**
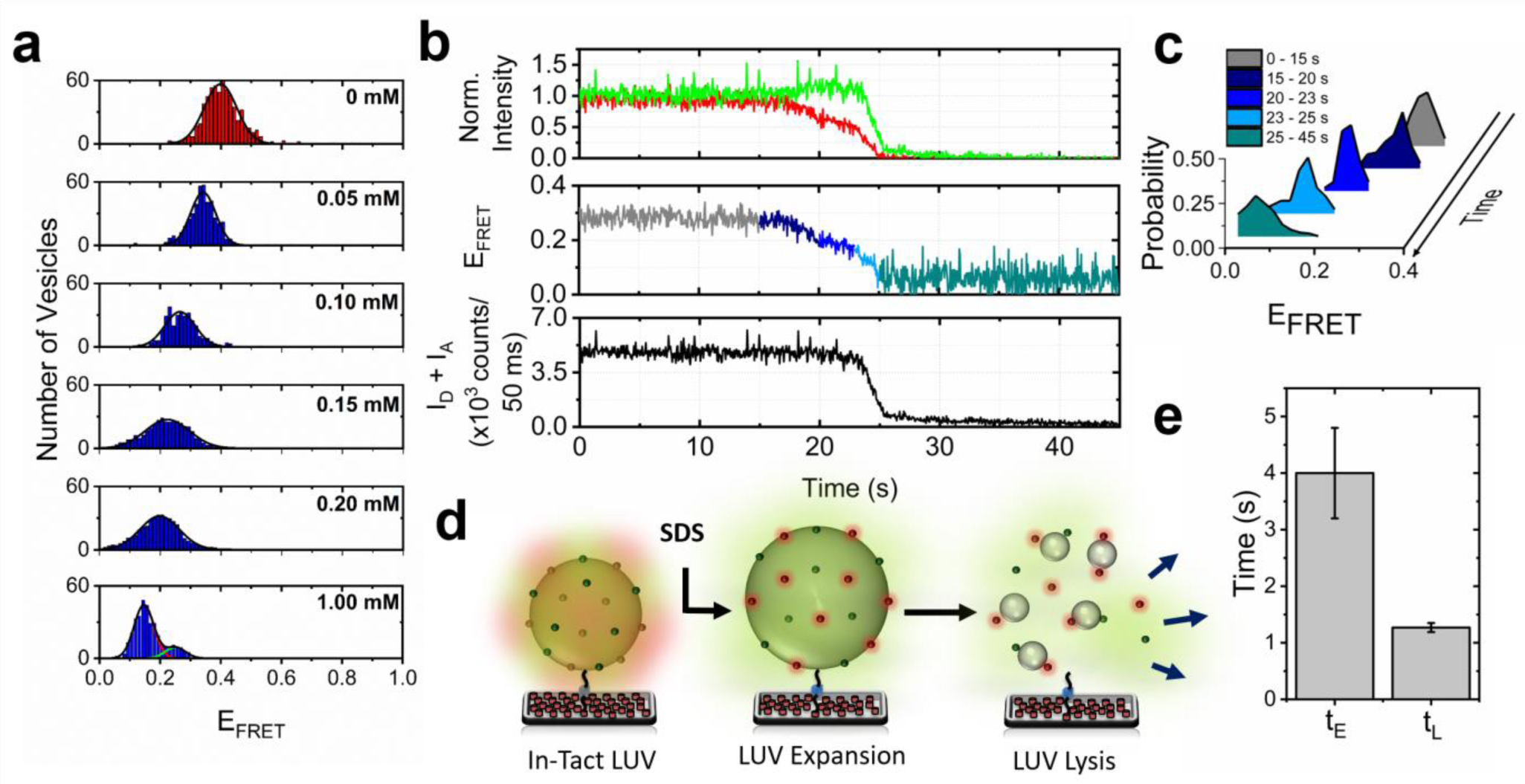
Real-time visualization of solubilization kinetics by svFRET. (a) E_FRET_ histograms obtained from immobilized LUVs after incubation with SDS at concentrations of 0 mM (N = 631), 0.05 mM (N = 438), 0.1 mM (N = 339), 0.15 mM (N = 400), 0.2 mM (N = 432) and 1 mM (N = 351). Solid lines represent Gaussian fits to the data. (b) Representative normalized variation in the fluorescence emission of Dil and DiD (top panel), the corresponding variation in FRET efficiency obtained before (< 15 s) and after (> 15 s) injection of 10 mM SDS (middle panel) and the corresponding sum of Dil and DiD fluorescence intensities (I_D_ + I_A_) (lower panel). (c) The corresponding relative FRET state occupancies observed over the 45 second measurement window. (d) Schematic illustration of the solubilization process. Injection of SDS leads to an initial LUV expansion event that precedes lysis. (e) Comparative bar plot summarizing the relative variation in t_E_ and t_L_ obtained after injection of 10 mM SDS. Error bars indicate the standard error of the mean (N = 30).

As we previously reported^[23]^, the different timescales and responses of both signals indicate that they represent different perturbations of the vesicle structure. First, the initial reduction in E_FRET_ with a largely unchanged total intensity indicates expansion of the LUV with little-to-no lipid loss, and second, a rapid decrease in total intensity while E_FRET_ remains invariant suggests a fast lysis step corresponding to release of lipids into solution (**Figure 3d**). The E_FRET_ plateau observed at ∼25 seconds when lipids are lost to solution (E_FRET_ 0.06 ± 0.01) represents an inter-dye distance of 8.4 ± 1.4 nm within the resultant micelle. Under these conditions, LUV expansion was found to occur with a half-life, t_E_, of 4.0 ± 0.8 seconds, followed by a faster lysis event with a half-life (T_L_) of 1.3 ± 0.1 seconds (**Figure 3e**). Since the expansion event reflects the initial interaction between SDS and the LUV, t_E_ can also be interpreted as the association and diffusion rate of SDS into the membrane.

The detection of a multi-step solubilization mechanism in LUVs agrees well with optical microscopy and phase contract measurements reported for giant unilamellar vesicles composed of 1-palmitoyl-2-oleoylglycero-3-phosphocholine (POPC) and DMPC lipids and with diameters > 1 μm^[21, 22]^. These experiments demonstrated that injection of SDS around the CMC causes an initial increase in the membrane surface area over the first several seconds, attributed to the homogeneous incorporation of SDS into the outer layer, which precedes membrane fracturing, the opening of transient nanopores and the ejection of mixed micelles. With increasing GUV size, the local density of SDS incorporation was found to differ across the membrane resulting in local instabilities. However, the microscopy techniques used in these studies only provide access to a cross-section of the GUVs and thus quantification of conformational changes across the entire three-dimensional volume is non-trivial. Nevertheless, the diffraction-limited LUVs and GUVs displayed common solubilization characteristics: comparable concentration requirements to achieve complete solubilization, and a vesicle swelling event that precedes a lysis step where micelles are released to the solution. Fluorescence microscopy experiments performed on POPC GUVs (10 – 20 μm) in the presence of the non-ionic detergent Triton X-100 (TX-100) at the CMC also showed an increase in surface area, attributed to the rapid insertion of TX-100 into the bilayer, prior to a gradual lysis step^[21, 32]^. Furthermore, when we applied svFRET technique to explore the effect of TX-100 on LUVs composed of POPC and 1-palmitoyl-2-oleoyl-sn-glycero-3-phospho-L-serine (POPS), the timescales associated with the expansion and lysis steps were found to be ∼5 s and ∼40 s, respectively^[23]^. Though it is possible that the kinetics vary as a function of the LUV composition, a similar expansion timescale was observed in the current work with SDS, while the lysis rate was an order of magnitude faster, consistent with previous studies investigating the interaction between anionic and non-ionic detergents and single cells^[33]^. Taken together, the svFRET studies on sub-micron and highly curved LUVs are complementary to phase-contrast, fluorescence and conventional optical microscopy experiments performed on GUVs, and both are consistent with a common general mechanism of detergent-induced membrane solubilization where a vesicle expansion event precedes lipid loss.

The ensemble FRET, DLS, AFM and svFRET data discussed so far afforded access to new structural and kinetic insights into the mechanism underpinning SDS-induced LUV solubilization. However, in order to quantify the extent of mass transfer at each stage, a complementary label-free QCM-D monitoring approach was employed. QCM-D has emerged as a powerful tool for monitoring vesicle deposition on solid surfaces^[34, 35]^, the formation of lipid bilayers^[34]^, vesicle fusion^[36]^ and protein-induced pore formation on supported bilayers^[37]^, but its utility in the context of probing detergent-LUV interactions has remained under-explored. Here, LUVs incorporating 1 % Biotin-PE were immobilized onto a QCM-D sensor using an identical immobilization strategy (**Figure S9**). Interactions between the immobilized LUVs and SDS were then monitored via real-time changes in the oscillation frequency and dissipation, reflecting the mass and viscoelasticity of the immobilized vesicles, respectively. As a solution of 0.6 mM SDS was flushed across the sensor surface, anti-correlative changes in both the frequency and dissipation traces were observed (**Figure 4a**), corresponding to a ∼17 % mass gain at the sensor surface, attributed to the initial deposition of SDS molecules onto the LUVs. This was followed by a non-destructive interaction that leads to a broad conformational change in in-tact vesicles. A considerable mass loss of 70 % was then observed via an increase in resonance frequency, consistent with immobilized material being released to solution (**Figure 4b**). The initial increase in energy dissipation observed as SDS entered the sensor surface relates to an increase in the surface viscoelasticity and LUV conformational changes arising from the deposition of SDS molecules. Following SDS deposition, the energy dissipation decreased substantially and closely matched that obtained from a control sensor lacking LUVs, indicating a decrease in viscoelasticity and loss of LUV surface material (**Figure 4a**). When a higher concentration (0.9 mM) was then flushed across the sensor, similar datasets were obtained (**Figure 4c, d**) pointing towards a mechanism through which mass is taken up by the LUVs prior to a conformational change and, at longer times, mass loss. Control experiments performed simultaneously indicated little interaction between a biotinylated-BSA-avidin coated sensor surface and SDS under the conditions tested, as indicated by minimal changes to the frequency and dissipation response after SDS injection across the sensors (**Figure 4a, c**). Taken together, this data set also supports a mechanism through which SDS accumulation on the highly-curved LUV surface precedes an expansion of the vesicle that in turn, precedes a lysis event. We note that while SDS may in addition promote membrane content leakage through pore formation^[38]^, this model is similar to that proposed for the non-ionic detergent TX-100 operating on LUVs of comparable size^[23]^ and supports a general mechanism of detergent-induced vesicle solubilization.

**Figure 4.**
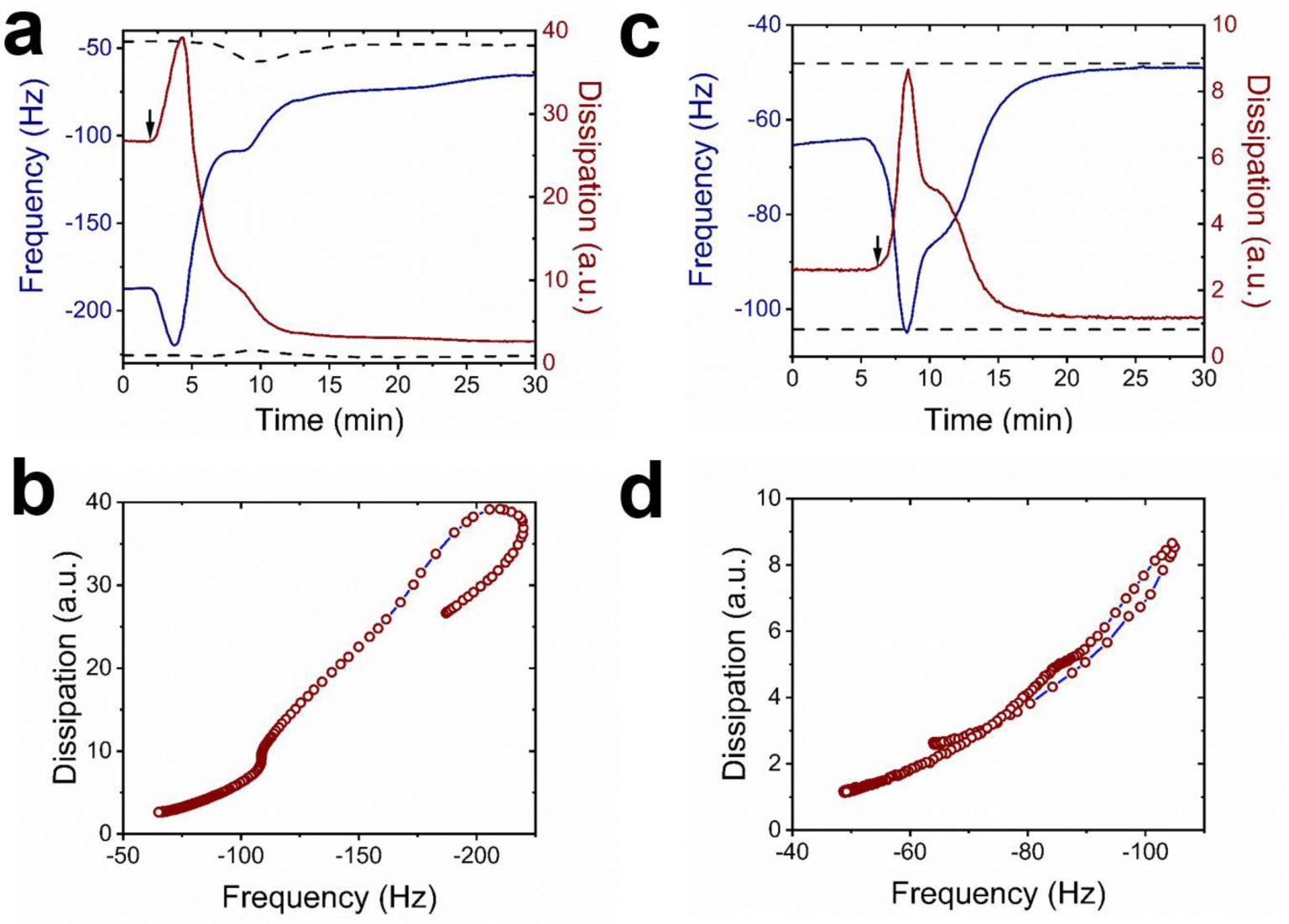
SDS induced vesicle solubilization monitored by QCM-D. (a) Representative variation in frequency (blue) and dissipation (red) of the 7^th^ overtone associated with surface immobilized LUVs in the presence of (a) 0.6 mM SDS and (c) 0.9 mM SDS. The dashed lines represent data collected from a control sensor pre-treated with biotinylated BSA and Avidin, but lacking LUVs. The arrows indicate the start-point of the solubilization process for each condition. The corresponding frequency versus dissipation plots observed during the interaction between surface immobilized vesicles and (b) 0.6 mM SDS and (d) 0.9 mM SDS.

## Conclusions

We have directly observed the solubilization of highly-curved LUVs in response to SDS using the combination of multiple advanced spectroscopic techniques. The collective data unambiguously separates each step of the SDS-induced solubilization pathway, and through the implementation of svFRET, kinetic parameters were assigned to each process without interference from vesicle fusion. We report the mechanism of SDS membrane solubilization as a sequence of events, whereby SDS deposition onto the LUV bilayer triggers expansion of in-tact LUVs that in-turn precedes rapid lipid loss. Exploring the organizational structure and real-time dynamics of controllable vesicles is especially appealing across the life sciences, not only because they are commonly used for a wide-variety of biotechnological applications, but also because critical trafficking pathways rely on the formation of highly-curved LUVs^[39, 40]^. We expect the presented approaches to be used for affording new access to previously intractable interactions between small molecule membrane disruptors and LUVs of varying size, to evaluate the role of lipid composition on the solubilization pathway, and to explore the interactions between a wide range of molecules that target, cross and disrupt the lipid membrane, such as those with important biomedical significance. We also anticipate that this work may lead to the quantification of the molecular mobility of membrane-integrated proteins relative to lipids observed in living cell systems^[41]^. Importantly, the observation of biphasic kinetics also has exciting implications for the adaptation and fine-tuning of detergent-based solubilization protocols.

## Methods

### Materials

DMPC and Biotin-PE phospholipids were purchased from Avanti Polar Lipids Inc. 1,1’-Dioctadecyl-3,3,3’,3’-Tetramethylindocarbocyanine Perchlorate (Dil) and 1,1’ -Dioctadecyl-3,3,3’,3’-Tet-ramethylindodicarbocyanine, 4 -Chlorobenzenesulfonate Salt) (DiD) were purchased from ThermoFisher Scientific. All phospholipid samples were used without additional purification and stored in chloroform at -20°C prior to use. Dil and DiD stock solutions were stored at 4°C prior to use. SDS was purchased from Sigma Aldrich and freshly suspended in 50 mM Tris (pH 8) buffer prior to each use.

### Preparation of large unilamellar vesicles

Mixtures of lipids and lipophilic dyes were homogeneously dispersed in chloroform, dried by nitrogen flow and stored under continuous vacuum pumping at 21°C for 5 hours. Phospholipid mixtures were subsequently re-suspended in buffer solution (50 mM Tris, pH 8) and mixed well by vortex. LUVs were prepared by the extrusion method^[42]^ in which they were passed through a 200 nm polycarbonate membrane filter. A molar ratio of 98.8: 1: 0.1: 0.1 DMPC: Biotin-PE: Dil: Dil was used for steady-state fluorescence, svFRET and AFM studies. For QCM-D work involving unlabeled LUVs, a molar ratio of 99: 1 DMPC: Biotin-PE was used. The size distribution of the prepared vesicles in solution was evaluated by dynamic light scattering using a Zetasizer μV molecular size detector (Malvern Instruments Ltd., UK).

### Steady-state fluorescence spectroscopy

Fluorescence emission spectra were acquired using a Horiba Fluoromax-4 fluorescence spectrophotometer. Spectra from Dil and DiD were recorded using an excitation wavelength of 532 nm. FRET efficiencies were approximated by the apparent FRET efficiency, E_FRET_ = (I_A_/[I_A_ + I_D_]), where I_A_ and I_D_ represent the fluorescence emission intensities of Dil and DiD, respectively. The FRET efficiency data shown in **Figure 1a** was fitted to a Hill model of the form, 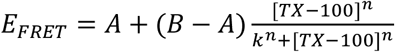, where A and B are the measured FRET efficiencies at the start and end of the titration, k is the half-maximal concentration constant and n is the Hill coefficient. The parameters of the fit shown in **Figure 1a** are A = 0.50 ± 0.02, B = 0.005 ± 0.001, k = 0.92 ± 0.16 mM and n = 1.25 ± 0.20 (χ^2^ = 0.99). Error bars represent the standard error of the mean from 3 individual experimental runs.

### Single-vesicle FRET Spectroscopy

Fluorescence emission at the donor and acceptor wavelengths were acquired from single vesicles by using a custom-built objective-type total internal reflection fluorescence microscope equipped with a continuous wave TEM_00_ 532 nm excitation line (Coherent, Obis). Pre-cleaned microscope slides were successively treated with 1 mg/mL bovine serum albumin (BSA), 0.1 mg/mL biotinylated BSA, and 0.2 mg/mL Avidin, before pM concentrations of freshly-prepared vesicles were added to the surface. Fluorescence movies were acquired with an integration time of 50 ms. The base buffer used for imaging was 50 mM Tris (pH 8), 6 % (w/v) glucose, 165 U/mL glucose oxidase, 2170 U/mL catalase and 2 mM trolox. Concentrations of SDS as specified in the main text were then included in the imaging buffer prior to being injected into the sample. Spatially-separated fluorescence images of donor and acceptor emission were collected using an oil immersion objective lens (NA = 1.49) and separated using a DualView emission splitter (Photometrics) containing a dichroic filter (T640LPXR, Chroma) and band pass filters (ET585/65M and ET700/75M, Chroma) on the donor and acceptor imaging paths. Dil and DiD emission intensities were collected in parallel using a cooled (−80°C) EMCCD camera (Andor iXON). The svFRET efficiency after background correction was evaluated via the apparent FRET efficiency, E_FRET_ = (I_A_/[I_A_ + I_D_]) ∼ R_o_ ^6^/([R_o_ ^6^+R^6^]), where I_A_ and I_D_ are the fluorescence intensities of the acceptor and donor, respectively, R_o_ is the Förster radius and R is the mean separation distance between the probes on each LUV. Half-lives were calculated by applying exponential fits to the E_FRET_ and [I_D_+I_A_] trajectories. Image processing was carried out using laboratory-written analysis routines developed in MATLAB 7. Excitation powers were measured immediately prior to laser light entering the back aperture of the objective lens.

### Quartz Crystal Microbalance with Dissipation (QCM-D) monitoring

QCM-D experiments were performed using a Q-sense E4 system (Biolin Scientific). SiO_2_-coated AT-cut quartz sensors (QSX 303, Biolin Scientific) were used, for which the fundamental frequency was 4.95 ± 0.05 MHz. The sensors were initially subjected to a 10 minute cleaning step by UV-ozone, prior to being sonicated in solutions of 2 % Hellmanex III and 2 x ultrapure Milli-Q water for 10 minutes. The sensors were then dried with N_2_ and placed under UV-ozone for a further 30 minutes. Each sensor was then immersed in 100% ethanol for 30 minutes and dried with N_2_ before installation in the flow modules. The QCM-D flow chambers were first flushed with ultrapure Milli-Q water for 1 hour, and then with 50 mM Tris buffer (pH 8) for 20 minutes before each measurement until a stable baseline was established (< 0.5 Hz shift over 10 min). The flow rate was kept constant at 20 μL/min. The sensor surfaces were then functionalized with 0.1 mg/mL biotinylated BSA and 1 mg/mL BSA then rinsed with 50 mM Tris buffer (pH 8.0) to remove unbound molecules. Thereafter, the sensor surface was flushed with 0.1 mg/mL Avidin solution in 50 mM Tris buffer (pH 8.0) for 20 min, followed by a rinse step with 50 mM Tris buffer (pH 8.0) for 20 min. Subsequently, vesicles coated with 1 % Biotin-PE were immobilized on the sensor surfaces by incubation with a 33 µg/mL vesicle solution for ∼80 minutes. SDS detergent solutions at the specified concentrations were then introduced into the QCM-D flow chambers. Changes in mass (Δm) were related to changes in frequency (Δf) via the Sauerbrey equation Δm = – (C · Δf)/n where n is the overtone number and C is a constant related to the properties of the quartz (17.7 ng Hz^-1^cm^-2^).

### Atomic Force Microscopy

Imaging of immobilized LUVs was conducted using a Bioscope Resolve Atomic Force Microscope (Bruker) in Fluid Tapping Mode. A silicon nitride cantilever (DNP-10 tips, Bruker) with nominal spring constant of 0.12 Nm^-1^ and resonant frequency of 23 kHz was used for all measurements. Typical scan sizes were 1.2 μm × 1.2 μm for probing multiple LUVs in 50 mM Tris buffer (pH 8). To evaluate LUV dimensions vesicles were sampled by taking multiple images per sample. Images were then selected to represent the average distribution, density and size of the sample. Structures were then sampled using the forbidden line unbiased counting rule in ImageJ. The mean caliper diameter at each LUV mid-height was measured both horizontally and vertically and the average of both measurements, d, was calculated.

## Author Contributions

S. D. Q. designed the research.

S. D. Q., J. J. –C., L. D., K. M., H. L., R. H. W. and A. B. performed the experiments.

S. D. Q., J. J. –C., S. J., M. C. L. and S. T. analyzed the data.

S. D. Q. wrote the manuscript with the help of all other authors.

## Additional Information

Figures S1-S9 and Tables S1 are found in the Supporting Information.

## Acknowledgements

We thank Dr. Daniella Barillà (University of York, UK) for use of the DLS instrumentation. We also thank the EPSRC (EP/P030017/1), BBSRC (BB/R001235/1) and Alzheimer’s Research UK (ARUK-RF2019A-001) for support.

## Data Availability Statement

The datasets generated during the current study are available from the corresponding author on reasonable request.

## References

1. Duquesne, K. and J.N. Sturgis, Membrane protein solubilization. Methods Mol Biol, 2010. 601: p. 205–17.

2. Prive, G.G., Detergents for the stabilization and crystallization of membrane proteins. Methods, 2007. 41(4): p. 388–397.

3. Zhou, Z.J., et al., Inactivation of viruses and bacteria on strawberries using a levulinic acid plus sodium dodecyl sulfate based sanitizer, taking sensorial and chemical food safety aspects into account. International Journal of Food Microbiology, 2017. 257: p. 176–182.

4. le Maire, M., P. Champeil, and J.V. Moller, Interaction of membrane proteins and lipids with solubilizing detergents. Biochim Biophys Acta, 2000. 1508(1-2): p. 86–111.

5. Lichtenberg, D., et al., Detergent solubilization of lipid bilayers: a balance of driving forces. Trends Biochem Sci, 2013. 38(2): p. 85–93.

6. Meister, A., A. Kerth, and A. Blume, Interaction of sodium dodecyl sulfate with dimyristoyl-sn-glycero-3-phosphocholine monolayers studied by infrared reflection absorption spectroscopy. A new method for the determination of surface partition coefficients. Journal of Physical Chemistry B, 2004. 108(24): p. 8371–8378.

7. Tan, A.M., et al., Thermodynamics of sodium dodecyl sulfate partitioning into lipid membranes. Biophysical Journal, 2002. 83(3): p. 1547–1556.

8. Apel-Paz, M., G.F. Doncel, and T.K. Vanderlick, Membrane perturbation by surfactant candidates for STD prevention. Langmuir, 2003. 19(3): p. 591–597.

9. Majhi, P.R. and A. Blume, Thermodynamic characterization of temperature-induced micellization and demicellization of detergents studied by differential scanning calorimetry. Langmuir, 2001. 17(13): p. 3844–3851.

10. Lopez, O., et al., Kinetic studies of liposome solubilization by sodium dodecyl sulfate based on a dynamic light scattering technique. Langmuir, 1998. 14(16): p. 4671–4674.

11. Hildebrand, A., et al., Temperature dependence of the interaction of cholate and deoxycholate with fluid model membranes and their solubilization into mixed micelles. Colloids and Surfaces B-Biointerfaces, 2003. 32(4): p. 335–351.

12. Helenius, A. and K. Simons, Solubilization of Membranes by Detergents. Biochimica Et Biophysica Acta, 1975. 415(1): p. 29–79.

13. Tan, A., et al., Thermodynamics of sodium dodecyl sulfate partitioning into lipid membranes. Biophys J, 2002. 83(3): p. 1547–56.

14. Mio, K. and C. Sato, Lipid environment of membrane proteins in cryo-EM based structural analysis. Biophys Rev, 2018. 10(2): p. 307–316.

15. Heerklotz, H., A.D. Tsamaloukas, and S. Keller, Monitoring detergent-mediated solubilization and reconstitution of lipid membranes by isothermal titration calorimetry. Nat Protoc, 2009. 4(5): p. 686–97.

16. Goni, F.M. and A. Alonso, Spectroscopic techniques in the study of membrane solubilization, reconstitution and permeabilization by detergents. Biochim Biophys Acta, 2000. 1508(1-2): p. 51–68.

17. Deo, N. and P. Somasundaran, Effects of sodium dodecyl sulfate on mixed liposome solubilization. Langmuir, 2003. 19(18): p. 7271–7275.

18. Bandyopadhyay, S., J.C. Shelley, and M.L. Klein, Molecular dynamics study of the effect of surfactant on a biomembrane. Journal of Physical Chemistry B, 2001. 105(25): p. 5979–5986.

19. Xu, B., et al., Molecular dynamics simulations of the effects of sodium dodecyl sulfate on lipid bilayer. Chinese Physics B, 2017. 26(3).

20. Yoshii, N. and S. Okazaki, A molecular dynamics study of structural stability of spherical SDS micelle as a function of its size. Chemical Physics Letters, 2006. 425(1-3): p. 58–61.

21. Sudbrack, T.P., et al., Observing the Solubilization of Lipid Bilayers by Detergents with Optical Microscopy of GUVs. Journal of Physical Chemistry B, 2011. 115(2): p. 269–277.

22. Igarashi, T., Y. Shoji, and K. Katayama, Anomalous Solubilization Behavior of Dimyristoylphosphatidylcholine Liposomes Induced by Sodium Dodecyl Sulfate Micelles. Analytical Sciences, 2012. 28(4): p. 345–350.

23. Dalgarno, P.A., et al., Unveiling the multi-step solubilization mechanism of sub-micron size vesicles by detergents. Scientific Reports, 2019. 9.

24. Nasr, G., et al., Liposomal membrane permeability assessment by fluorescence techniques: Main permeabilizing agents, applications and challenges. Int J Pharm, 2020. 580: p. 119198.

25. Elsayed, M.M.A., M.M. Ibrahim, and G. Cevc, The effect of membrane softeners on rigidity of lipid vesicle bilayers: Derivation from vesicle size changes. Chemistry and Physics of Lipids, 2018. 210: p. 98–108.

26. Niroomand, H., et al., Lipid-Detergent Phase Transitions During Detergent-Mediated Liposome Solubilization. Journal of Membrane Biology, 2016. 249(4): p. 523–538.

27. Vanni, S., et al., A sub-nanometre view of how membrane curvature and composition modulate lipid packing and protein recruitment. Nature Communications, 2014. 5.

28. Sebaihi, N., et al., Dimensional characterization of extracellular vesicles using atomic force microscopy. Measurement Science and Technology, 2017. 28(3).

29. Woo, J., S. Sharma, and J. Gimzewski, The Role of Isolation Methods on a Nanoscale Surface Structure and its Effect on the Size of Exosomes. J Circ Biomark, 2016. 5: p. 11.

30. Sharma, S., et al., Nanofilaments on glioblastoma exosomes revealed by peak force microscopy. J R Soc Interface, 2014. 11(92): p. 20131150.

31. Parisse, P., et al., Atomic force microscopy analysis of extracellular vesicles. Eur Biophys J, 2017. 46(8): p. 813–820.

32. Mattei, B., A.D. Franca, and K.A. Riske, Solubilization of binary lipid mixtures by the detergent Triton X-100: the role of cholesterol. Langmuir, 2015. 31(1): p. 378–86.

33. Brown, R.B. and J. Audet, Current techniques for single-cell lysis. J R Soc Interface, 2008. 5 Suppl 2: p. S131–8.

34. Lind, T.K., M. Cardenas, and H.P. Wacklin, Formation of supported lipid bilayers by vesicle fusion: effect of deposition temperature. Langmuir, 2014. 30(25): p. 7259–63.

35. Richter, R., A. Mukhopadhyay, and A. Brisson, Pathways of lipid vesicle deposition on solid surfaces: a combined QCM-D and AFM study. Biophys J, 2003. 85(5): p. 3035–47.

36. Morigaki, K. and K. Tawa, Vesicle fusion studied by surface plasmon resonance and surface plasmon fluorescence spectroscopy. Biophys J, 2006. 91(4): p. 1380–7.

37. Briand, E., et al., Combined QCM-D and EIS study of supported lipid bilayer formation and interaction with pore-forming peptides. Analyst, 2010. 135(2): p. 343–50.

38. Keller, S., et al., Thermodynamics of lipid membrane solubilization by sodium dodecyl sulfate. Biophysical Journal, 2006. 90(12): p. 4509–4521.

39. Shibata, Y., et al., Mechanisms shaping the membranes of cellular organelles. Annu Rev Cell Dev Biol, 2009. 25: p. 329–54.

40. Antonny, B., Mechanisms of Membrane Curvature Sensing. Annual Review of Biochemistry, Vol 80, 2011. 80: p. 101–123.

41. Nenninger, A., et al., Independent mobility of proteins and lipids in the plasma membrane of Escherichia coli. Mol Microbiol, 2014. 92(5): p. 1142–53.

42. Olson, F., et al., Preparation of Liposomes of Defined Size Distribution by Extrusion through Polycarbonate Membranes. Biochimica Et Biophysica Acta, 1979. 557(1): p. 9–23.

