## Supplementary Information for "The Mechanism of Vesicle Solubilization by the Detergent Sodium Dodecyl Sulfate"

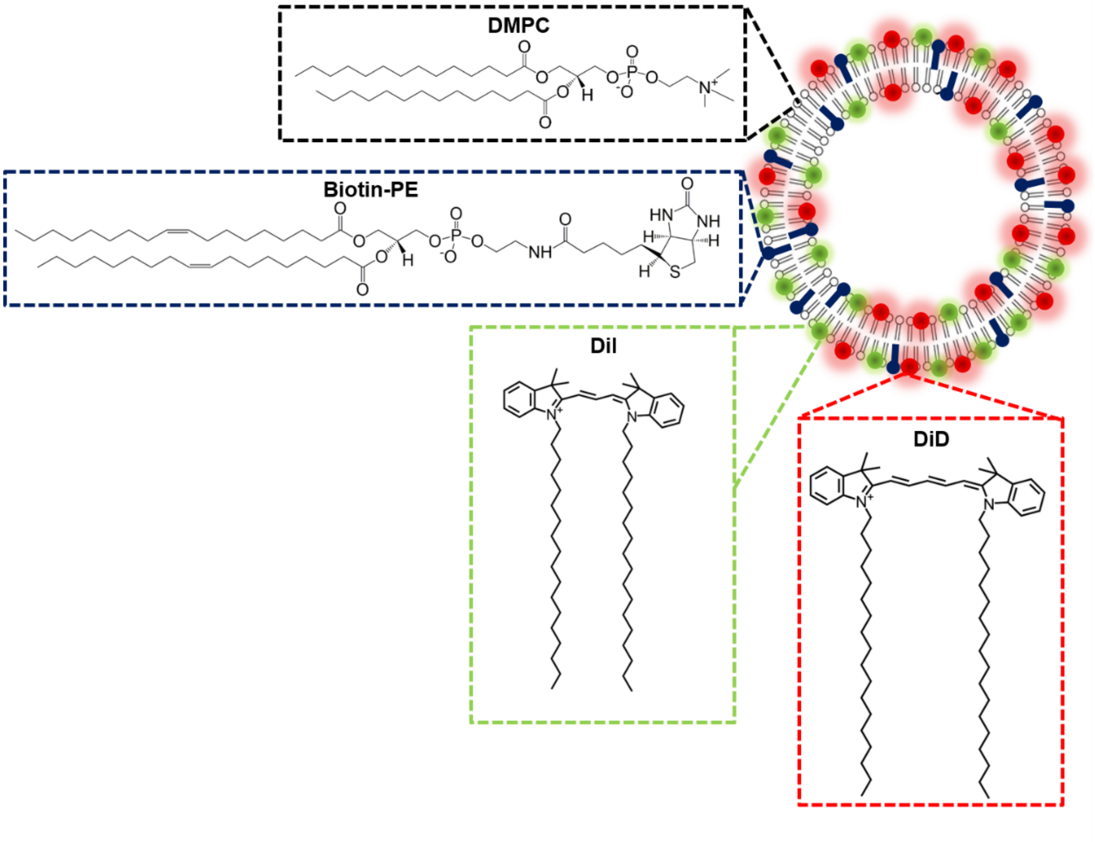


**Figure S1.** Schematic representation of LUVs composed of 98.8 % DMPC, 1 % Biotin-PE, 0.1 % DiI and 0.1 % DiD. Insets: the chemical structures of DMPC (black), Biotin-PE (blue), DiI (green) and DiD (red) are shown.

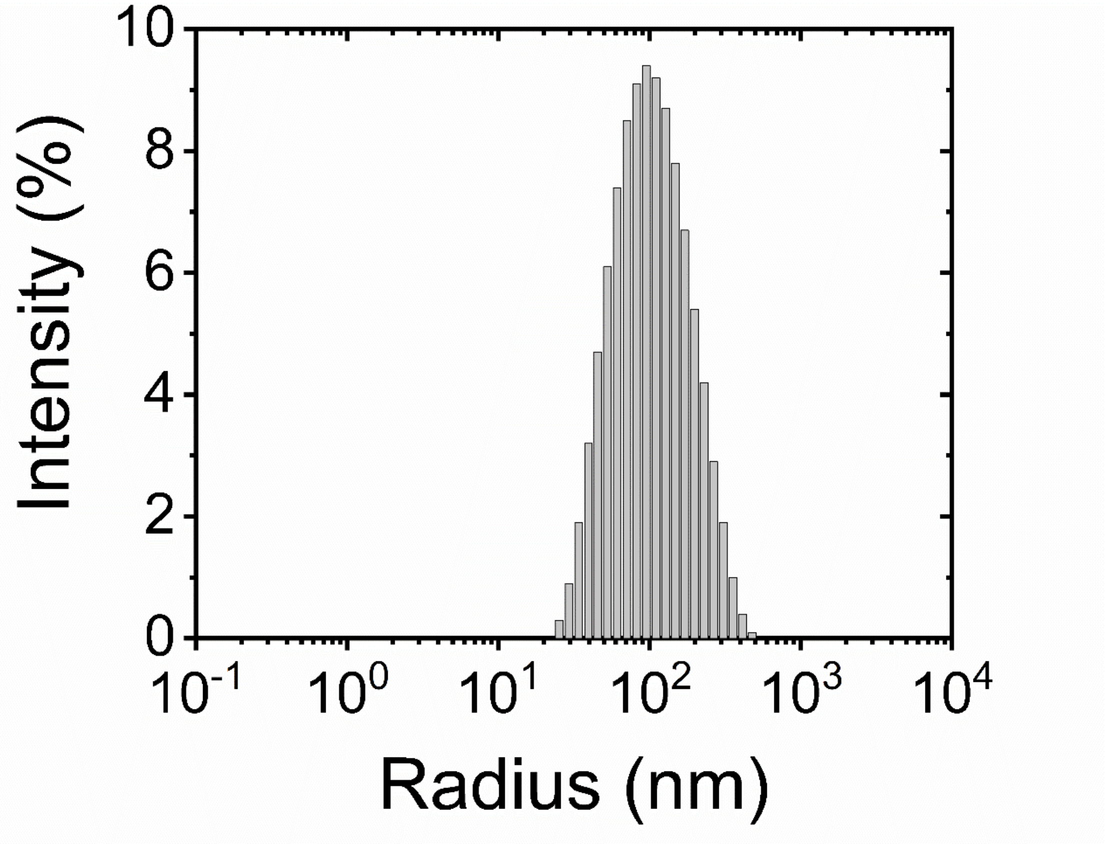


**Figure S2**. Semi-log plot of LUV hydrodynamic radius as a function of scattering intensity, as obtained by dynamic light scattering. Data shown represents the mean obtained from 3 experimental runs.

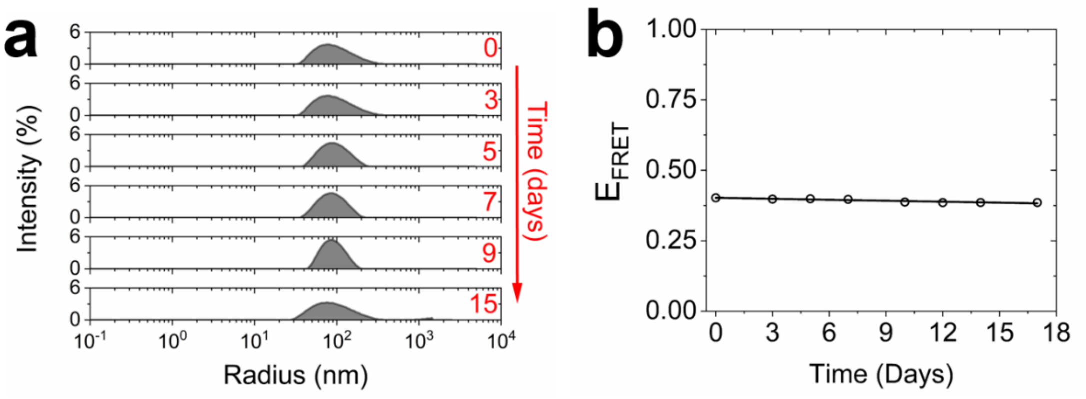


**Figure S3**. **LUV stability monitored by DLS and steady-state FRET spectroscopy.** (a) Semi-log plot of LUV hydrodynamic radius as a function of scattering intensity and (b) ensemble $E_{FRET}$ obtained from LUVs in 50 mM Tris (pH 8.0) buffer as a function of time. LUVs were freshly prepared by the extrusion method on day 0. Data shown represents the mean values obtained from 3 experimental runs.

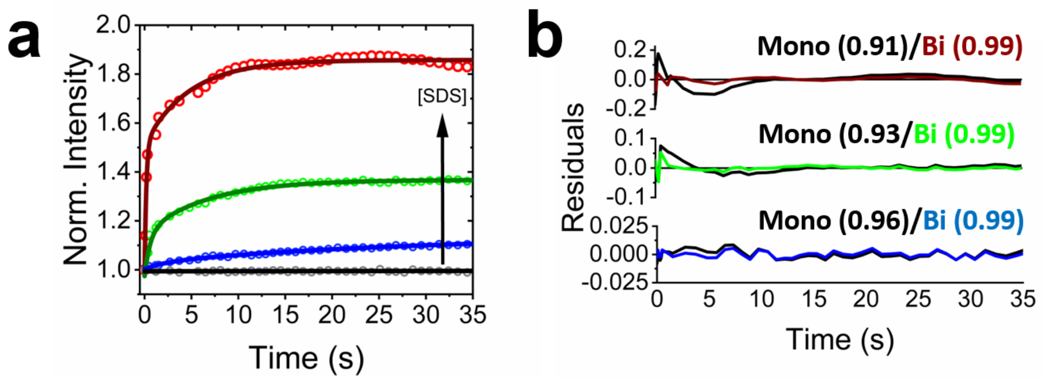


**Figure S4. Ensemble kinetics of SDS solubilization.** (a) Normalized representative variation in the fluorescence emission intensity of DiI as a function of time with no SDS (grey) and following injection of 1 mM (blue), 3 mM (green) and 4 mM (red) mM SDS. The solid black line represents a linear fit ($R^2$ = 0.98) and the blue, green and red solid lines represent bi-exponential fits. (b) Results obtained for each growth after fitting to a mono- or biexponential function are also shown. Numbers shown in brackets represent $R^2$ values.

**Table S1** Pre-exponential factors, time constants and fitting errors associated with the time-dependent fluorescence enhancement of DiI emission as a function of SDS. Experimental conditions: 50 mM Tris, pH 8, 21°C. Vesicles were composed of 98.8 % DMPC, 1 % Biotin-PE, 0.1 % DiI and 0.1 % DiD.

| | **1 mM** | **3 mM** | **4 mM** |
|---|---|---|---|
| **$a_1$** | 0.03 ± 0.01 | 0.17 ± 0.01 | 0.53 ± 0.03 |
| **$\tau_1$ (s)** | 2.13 ± 0.08 | 0.58 ± 0.13 | 0.18 ± 0.02 |
| **$a_2$** | 0.11 ± 0.01 | 0.22 ± 0.02 | 0.34 ± 0.02 |
| **$\tau_2$ (s)** | 28.7 ± 7.7 | 6.67 ± 0.86 | 4.77 ± 0.51 |
| **$R^2$** | 0.992 | 0.987 | 0.988 |
| ***$\tau_{av}^*$ (s)*** | ***28.2 ± 7.6*** | ***6.28 ± 0.81*** | ***4.51 ± 0.48*** |

* Amplitude weighted average time constants, $\tau_{av}$, were calculated according to the equation $\tau_{av} = \sum_{i=1}^{N} a_i \tau_i^2 / \sum_{i=1}^{N} a_i \tau_i$ where $a_i$ represents the pre-exponential factor and $\tau_i$ is the associated time constant.

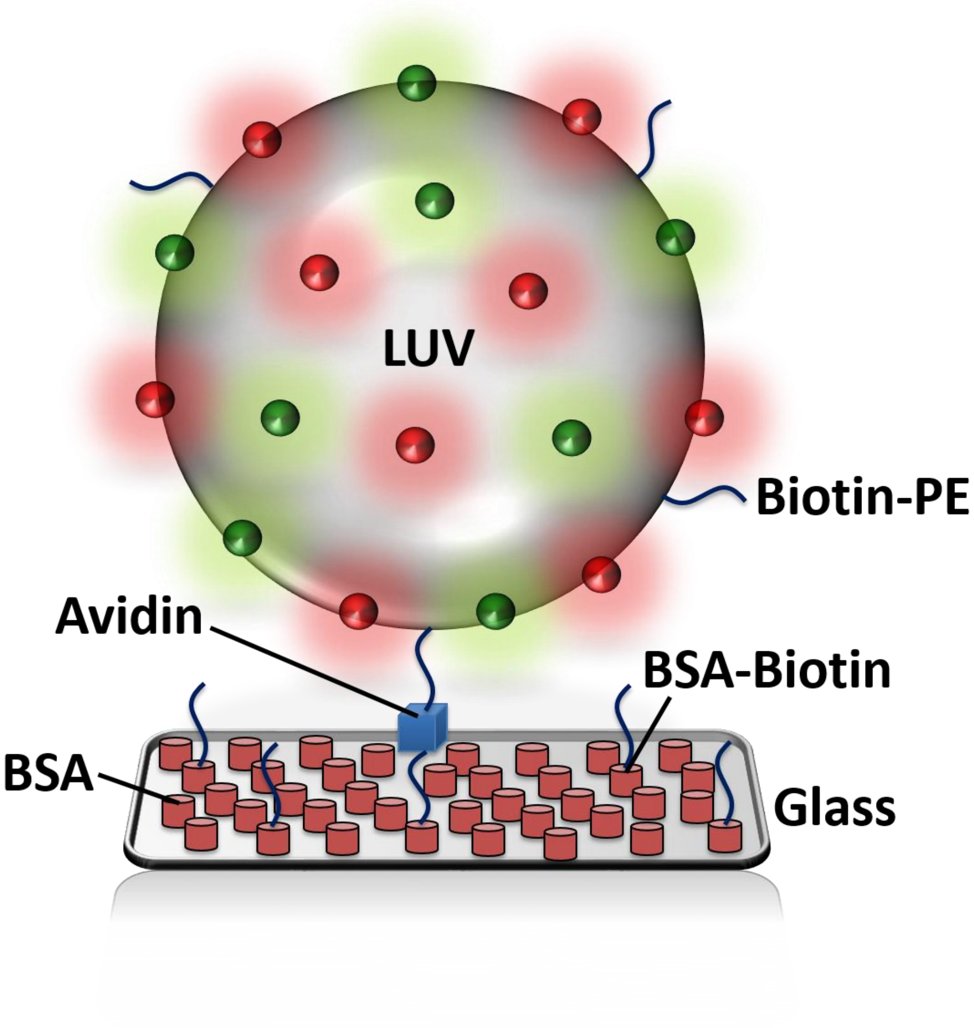


**Figure S5.** Schematic illustration of the immobilization scheme used to couple biotinylated vesicles to a glass substrate. Here, a glass substrate coated in 99 % BSA and 1 % biotinylated BSA is used to anchor biotinylated LUVs to the surface via Avidin.

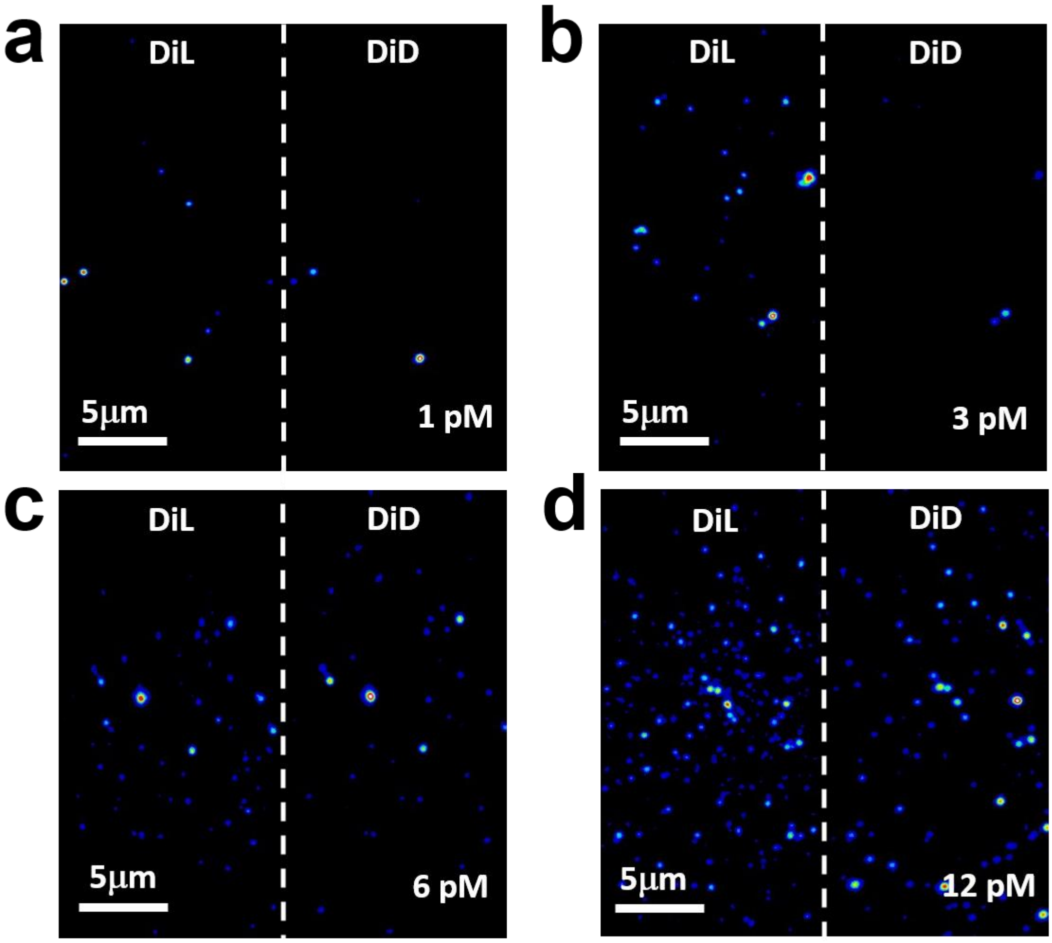


**Figure S6.** Representative total internal reflection fluorescence (TIRF) images of immobilized LUVs composed of 98.8 % DMPC, 1 % Biotin-PE, 0.1 % Dil and 0.1 % DiD at LUV concentrations of (a) 1 pM, (b) 3 pM, (c) 6 pM and (d) 12 pM, with $\lambda_{ex}$ = 532 nm.

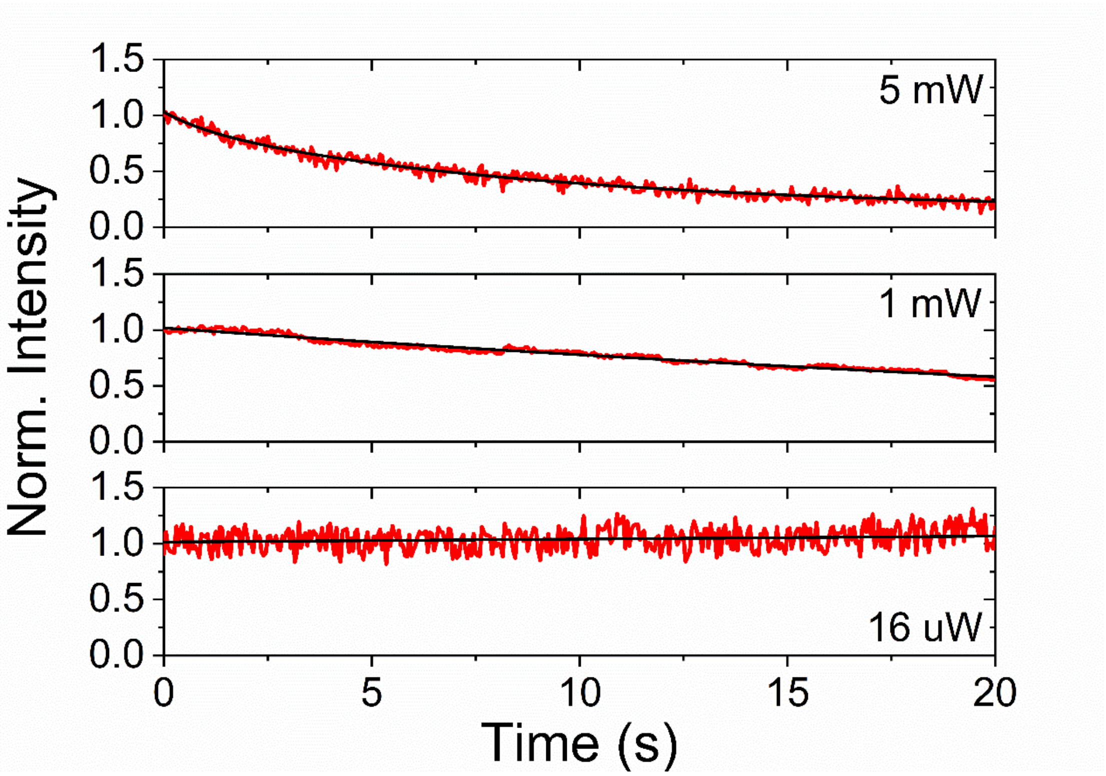


**Figure S7.** Representative normalized fluorescence trajectories of DiD obtained from single DiI/DiD labelled LUVs as a function of excitation intensity with $\lambda_{ex}$ = 532 nm.

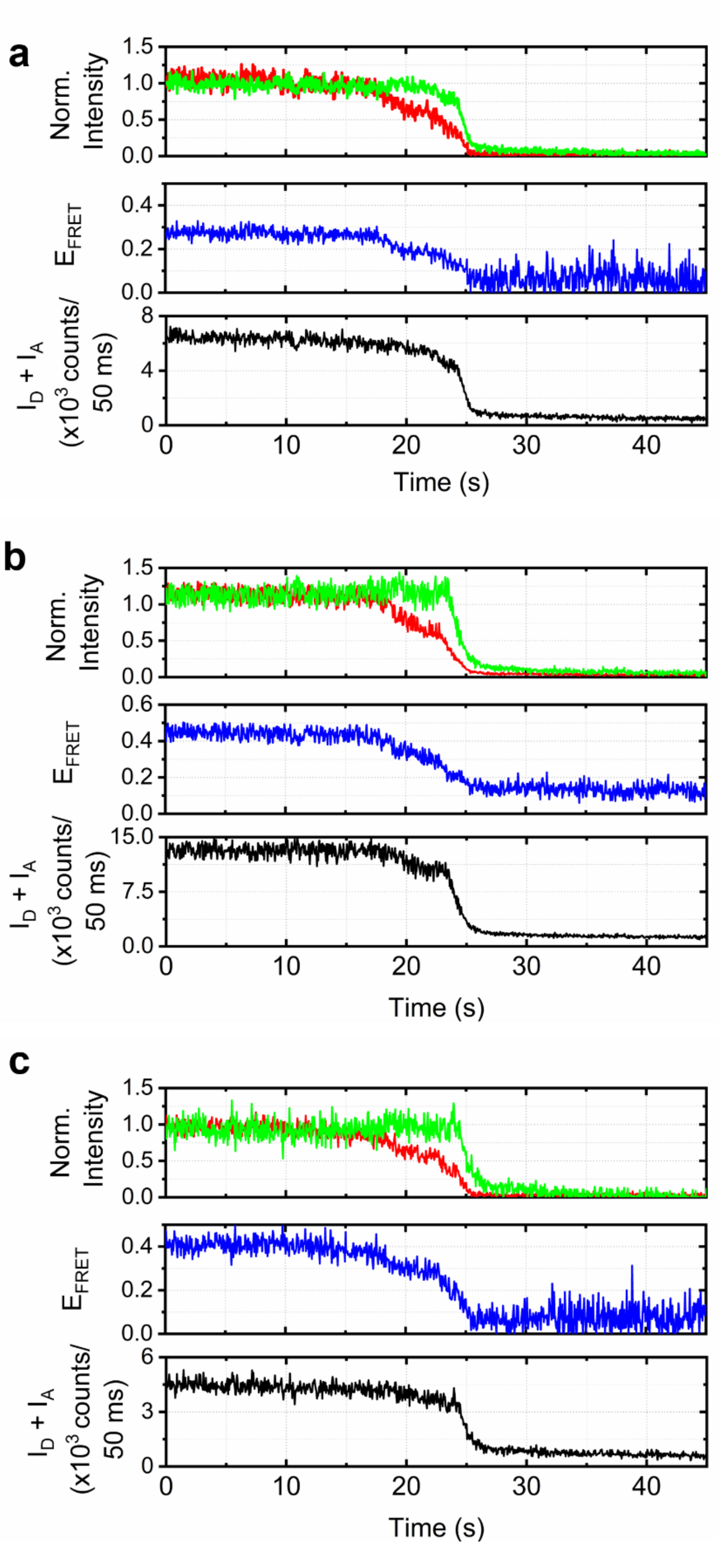


**Figure S8.** (a-c) Representative normalized fluorescence trajectories (top panel), corresponding FRET efficiency (middle panel) and total intensity (lower panel) obtained from single DiI (green) and DiD (red) labelled LUVs before and after injection of 10 mM SDS, with $\lambda_{ex}$ = 532 nm an excitation power of 16 $\mu$W.

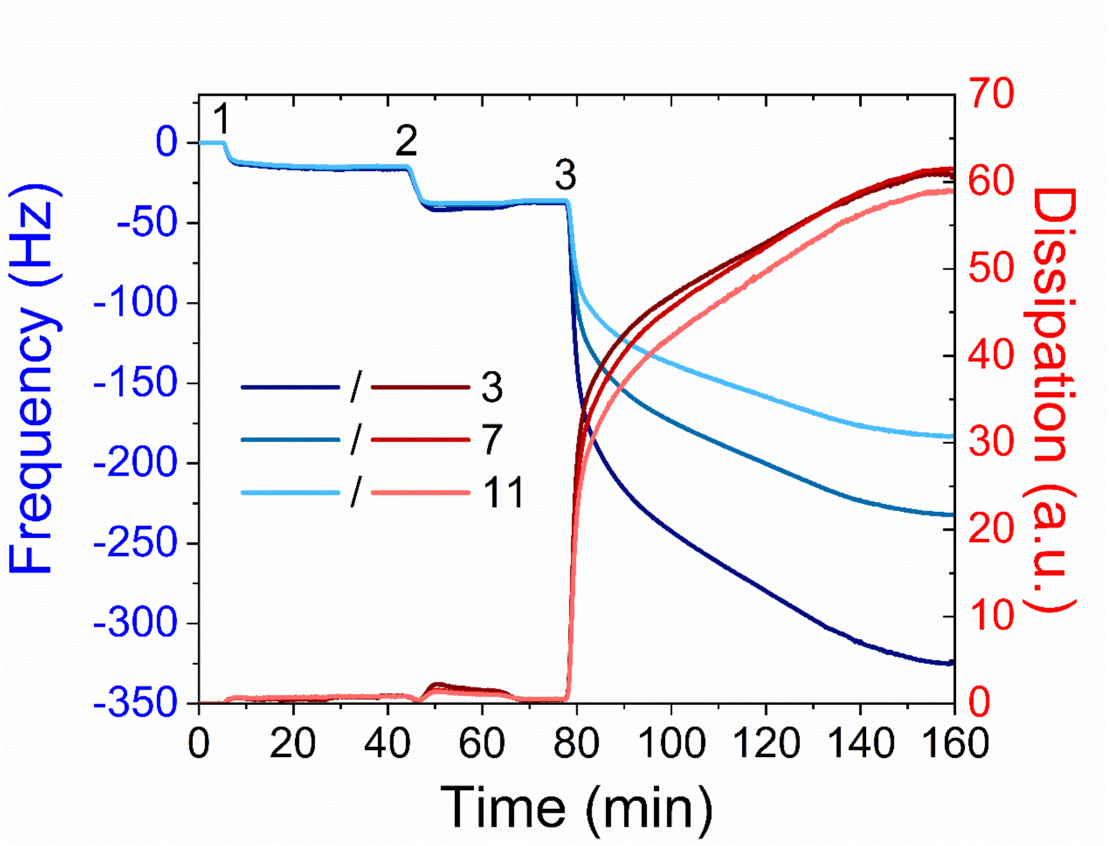


**Figure S9**. **Real-time LUV immobilization monitored by QCM-D.** Variation in frequency (blue curves) and dissipation (red curves) associated with the 3$^{rd}$, 7$^{th}$ and 11$^{th}$ overtone as BSA-Biotin (1), avidin (2) and LUVs (3) were flushed across the sensor surface sequentially. The sensor surface was washed with 50 mM Tris (pH 8) buffer between each step.
